# Limits for resolving tandem mass tag reporter ions with identical integer mass using phase constrained spectrum deconvolution

**DOI:** 10.1101/332668

**Authors:** Christian D. Kelstrup, Konstantin Aizikov, Tanveer S. Batth, Arne Kreutzman, Dmitry Grinfeld, Oliver Lange, Daniel Mourad, Alexander Makarov, Jesper V. Olsen

## Abstract

A popular method for peptide quantification relies on isobaric labeling such as tandem mass tags (TMT) which enables multiplexed proteome analyses. Quantification is achieved by reporter ions generated by fragmentation in a tandem mass spectrometer. However, with higher degrees of multiplexing, the smaller mass differences between the reporter ions increase the mass resolving power requirements. This contrasts with faster peptide sequencing capabilities enabled by lowered mass resolution on Orbitrap instruments. It is therefore important to determine the mass resolution limits for highly multiplexed quantification when maximizing proteome depth. Here we defined the lower boundaries for resolving TMT reporter ions with 0.0063 Da mass differences using an ultra-high-field Orbitrap mass spectrometer. We found the optimal method depends on the relative ratio between closely spaced reporter ions and that 64 ms transient acquisition time provided sufficient resolving power for separating TMT reporter ions with absolute ratio changes up to 16-fold. Furthermore, a 32 ms transient processed with phase-constrained spectrum deconvolution provides >50% more identifications with >99% quantified, but with a slight loss in quantification precision and accuracy. These findings should guide decisions on what Orbitrap resolution settings to use in future proteomics experiments relying on TMT reporter ion quantification with identical integer masses.

## INTRODUCTION

Quantitative shotgun proteomics deals with large-scale measurements of peptide abundances using liquid chromatography (LC) coupled to mass spectrometry (MS), where quantification is achieved through either stable isotope dilution or label free quantification (LFQ) methods. A prominent choice for isotopic labeling is the isobaric labeling of peptides using TMT reagent sets.^1,2^ This chemical labeling approach enables multiplexing via a stable heavy isotopically coded tag that consists of an amine-reactive group, a balancer group and a reporter ion group. The differentially TMT labeled peptides are indistinguishable in mass but fragmentation enables relative quantification using the differential reporter ions that are released upon collisionally-induced dissociation (CID). While the balancer group can also be used for quantification^3,4^, the most commonly used approach is to use the characteristic reporter ions. These were initially limited in multiplexing to four-or six-plex, but with the higher resolution tandem MS (MS/MS) instrumentation becoming readily available, the mass differences below integer masses can be resolved and therefore a higher sample multiplexing can be achieved.^5,6^ However, the required mass resolution is demanding even for modern MS instruments associated with the overhead costs related to the speed of acquisition as resolving power of Fourier transform (FT)-MS scales with acquisition time.

An MS instrument is a central part of the general shotgun proteomics workflow and a key element for fast analysis of highly multiplexed TMT labeled samples. One of the most popular mass analyzers for proteomics is the Orbitrap^™^ mass analyzer, which belongs to the FT-MS family of instruments.^7,8^ Enhanced FT (eFT^™^) calculation^9,10^ has increased the obtained resolution for the same transient signal and although it is superior to conventional (magnitude) FT, the enhanced method has a fundamental limit in achievable resolution imposed by Fourier uncertainty. In recent years novel super-FT resolving methods has demonstrated the ability to resolve isobaric TMT reporter ions on FT-MS transients substantially shorter than otherwise would be required by FT based signal processing techniques.^11,12^ Particularly, a new computational approach termed the phase-constrained spectrum deconvolution method (ΦSDM) promises further improvements in spectral quality and MS acquisition rate albeit at an additional computational cost.^13^

Here, we experimentally investigate the lower limit of the mass resolution required to resolve closely spaced reporter ions with identical integer masses. We further describe an implementation of ΦSDM FT-MS that enables higher resolution for TMT reporter ions with short acquisition transients in real time as they are kept within the limitations of the existing hardware. This is accomplished by applying ΦSDM only in a very narrow frequency/mass band containing the reporter ions.

The focus is kept on the resolution requirements for TMT10-plex which are here seen as independent to a broader discussion of general limitations of the TMT labeling strategy. These have been described elsewhere in great detail and include labeling challenges, accuracy challenges due to co-fragmentation of impure precursors and altered gas phase fragmentation and charge state distributions.^14–17^ The narrow aim of the present work has been to describe the trade-offs with fast acquisition methods for quantitative proteomics with TMT10-plex labeling.

## MATERIALS AND METHODS

### Instrument modifications

Transients were processed directly on the instrument computer of the commercial Q Exactive^™^ HF-X mass spectrometer (Thermo Fisher Scientific, Bremen, Germany) with a research-grade data analysis software with in-house implementation of the ΦSDM algorithm. eFT processing was conducted with the default settings at MS/MS transient lengths of 16, 32, 64, 96, and 128 ms. To ensure the real time computation of ΦSDM, it was calculated in 0.22 Da wide spectral windows centered at the reporter ion regions of a TMT 10-plex at 126, 127, 128, 129, 130, and 131 Da. The phase was constrained to within a ±5° cone. The number of iterations was set to 50. Frequency grid refinement was set to 16.

### Cell culture

Immortalized human epithelial cervix carcinoma adherent cells (HeLa) were grown in 15 centimeter dishes in DMEM (Gibco, Waltham, USA) media containing 2 mM L-glutamine supplemented with 10% fetal bovine serum (Gibco, Waltham, USA), 100 U/ml penicillin (Life Technologies, Carlsbad, USA), 100 μg/ml streptomycin in 37°C incubator supplemented with 5% CO_2_. Growing cells were allowed to obtain between 80-90% confluency prior to harvesting.

### Sample preparation and trypsin digestion

Media was removed from the plates and the cells washed twice with cold phosphate containing buffer (1x PBS). Cells were rapidly lysed and cysteines were reduced and alkylated in a single step as previously described.^18^ Briefly, 4ml of boiling 6M guanidine hydrochloride (Gnd-HCl) containing 10 mM chloroacetamide and 10 mM tris(2-carboxyethyl)phosphine was added directly to the plates and the cells were manually collected by scraping. Lysis buffer containing cells were boiled for an additional 10 minutes at 99°C followed by sonication (Vibra-Cell VCX130, Sonics, Newtown, CT, USA) for 2 minutes with pulses of 1 s on and 1 s off at 50% amplitude. Protein concentration was determined by Bradford assay (Bio-Rad, Hercules, USA).

Lys-C protease (Wako Chemicals, Richmond, USA) was added at a ratio of 1:100 (w/w) and digested for 2 hours at 37 °C. The concentration of Gnd-HCl was reduced to less than 1M with 25mM ammonium bicarbonate prior to the addition of trypsin (Life Technologies, Carlsbad, USA) at a ratio of 1:50 and allowed to digest overnight at 37 °C. After digestion, the sample was acidified using trifluoroacetic acid (TFA) to a 1% final concentration in order to quench protease activity followed by centrifugation for 5 minutes at 4000 g to remove residual cell debris. Supernatant was cleared of all salts/buffers and tryptic peptides were purified using solid phase extraction on C_18_ Sep-Pak (Waters Corporation, Milford, USA) by gravity. Peptides were eluted with 50% acetonitrile (ACN), 0.1% formic acid (FA). Peptides were dried and ACN was removed from the peptide containing sample by SpeedVac (Eppendorf, Germany) at 45°C. Peptide concentration was measured using a NanoDrop^™^ 2000 spectrometer (Thermo Fisher Scientific, Waltham, USA).

### TMT labeling

Peptides were labeled using TMT-10plex isobaric tags (Thermo Fisher Scientific, San Jose, USA) according to manufacturer’s instructions. Briefly, peptides were dissolved in 40% ACN containing 40 mM HEPES buffer at pH 8.5. TMT reagent dissolved in neat ACN was added to peptide samples and the reaction was carried out at room temperature for 1 hour. Reaction was quenched with the addition of 1% hydroxylamine for 15 minutes. TMT-plex 1-10 labeled samples were mixed into one and acidified to 1% TFA. ACN of labeled mixed samples was evaporated by speedvac and additional buffers and excess TMT reagent was removed using C_18_ solid phase extraction as described above. Peptide concentration of labeled samples was again measured using the NanoDrop 2000 spectrometer.

### Nanoflow LC/MS/MS

The nano LC/MS/MS analysis was performed as previously explained with few alterations.^19^ Briefly, the peptide solution was adjusted in volume to 0.1 µg/µl and kept in loading buffer (5% ACN and 0.1% TFA) prior to autosampling. The EASY-nLC^™^ 1200 system (Thermo Fisher Scientific, San Jose, USA) was equipped with an integrated column oven (PRSO-V1, Sonation GmbH, Biberach, Germany) maintaining temperature at 40 °C for the in-house packed 15 cm, 75 µm ID analytical capillary column with 1.9 μm Reprosil-Pur C_18_ beads (Dr. Maisch, Ammerbuch, Germany) which was interfaced online with the mass spectrometer. FA 0.1% was used to buffer the pH in the two running buffers. The gradient went from 8% to 24% ACN in 12.5 minutes followed by 24% to 36% in 2.5 minutes. This was followed by a washout by a 0.5 minute increase to 64% ACN which was kept for 4.5 minutes. Flow rate was kept at 350 nL/minute. Re-equilibration was done in parallel with 2 µl sample pickup and prior to loading with a minimum requirement of 0.5 µL 0.1% FA buffer at a pressure of 800 bar.

The Q Exactive HF-X instrument was configured as described above with custom Tune (version 2.9) instrument control software that enabled a Tune toggle for ΦSDM for TMT10-plex. Spray voltage was set to 2 kV in positive polarity, funnel RF level at 40, and heated ion transfer tube temperature at 275 °C. The methods employed were Full MS/ DD-MS/MS. Full scan resolutions were set to 60,000 at m/z 200 and full MS scan target was 3E6 with an IT of 45 ms. Mass range was set to 350-1400. Target value for fragment scans was set at 1E5, and intensity threshold was kept at 1E5. Isolation width was set at 0.8 m/z. Dynamic exclusion was enabled and set at 30 s. A fixed first mass of 100 m/z was used. Normalized collision energy was set at 33%. Peptide match was set to off, and isotope exclusion was on. Charge state exclusion rejected ions having unassigned charge states or that were singly charged or had a charge state above 5. Full MS data were acquired in the profile mode with fragment scans recorded in the centroid mode. For each of the MS/MS transient lengths, specific settings were set to keep parallel acquisition optimal and MS cycle time approximately constant. The MS/MS 7500 resolution or 16ms transient was combined with a loop count of 40 and an injection time of 11ms. The 15,000 resolution or 32ms transient was combined with a loop count of 28 and an injection time of 22 ms. The 30,000 resolution or 64ms transient had a loop count at 14 and an injection time of 54 ms. The 60,000 or 128 ms transient had a loop count of 7 and an injection time of 118 ms. A separate control experiment was performed on all samples just after the primary experiment where all instrument methods were also measured with a constant 86ms injection time and a loop count of 10.

### Data analysis

All raw files were analyzed in one combined analysis using the MaxQuant software suite (version 1.6.1.2) with the integrated Andromeda search engine, where each raw file was configured as a separate experiment.^20^ For the search, fixed modifications were set to carbamylation of cysteines and TMT-10 were specified as label on N-terminal and lysines residues with mass tolerance set to 0.003 Da. Variable modifications were set to oxidation of methionines, protein N-terminal acetylation, deamidation of asparagine or glutamine, and pyro-glutamate formation from N-terminal glutamine. The database was the UniProt human reference proteome release 2018_02 without isoforms containing 21,010 protein sequences. In addition, the included contaminant database from MaxQuant contains enzymes and common contaminants. FDR was set to 1% on the PSM, site, and protein level. Otherwise default values were used.

All data were filtered to only focus on identified spectra. Reading of intensity-to-noise values was performed with the raxport command line tool (freely available at http://code.google.com/p/raxport/) together with custom scripts in Perl to filter the large lists. Subsequent data analysis was performed in R.^21^

The mass spectrometry proteomics data have been deposited to the ProteomeXchange Consortium^22^ via the PRIDE partner repository with the data set identifier PXD009821. Reviewer account details:

Username

Password: fPrhmkbF

## RESULTS AND DISCUSSION

TMT10-plex quantification relies on resolution and accurate measurements of the reporter ions with mass differences as small as 3.6 mDa, which places a high demand on the mass resolving power of the MS/MS measurements. It is known, however, that resolution and speed are a trade-off for FT-MS instruments and increases in acquisition speed has been shown to improve throughput and increase proteome coverage depth.^19,23^ It is therefore important to uncover the lower limits of the resolution requirements for resolving TMT reporter ions with identical integer masses, and thereby establish resolution settings that are sufficient to achieve accurate quantification, as well as settings at which no practical benefits with higher resolution is observed. Further, TMT quantification is expected to benefit from new signal processing methods targeted to improve the obtained resolution.

Traditionally, separating closely spaced MS peaks required them to be baseline resolved or have a low degree of overlap known as the ‘10% valley resolution definition’.^24^ This criterion mainly deals with peaks of similar height and largely ignores that it is more difficult to separate peaks that have ratios of 1:4 or higher. This is easily demonstrated through modeling where resolution needs to be higher in order to separate peaks of different intensities (Figure 1a). Due to this effect, an insufficient resolving power results in closely spaced peaks appearing either merged or having their intensities corrupted by the FT interferences.^25^ For high abundance ratios, the smaller peak might simply disappear in the noise band and be reported as a missing data-point. Missing data-points are usually not a problem in the TMT-based quantification when sufficient MS/MS resolution settings are used. However, for the low resolution settings, missing data-points start to appear, which can be used as a simple measure for how well the quantitative method separates closely spaced peaks. Other possible indicators can be the mass accuracy, which might be affected if the peaks merge, or quantitative descriptive parameters such as accuracy and precision. These considerations form the basis for the current report.

**Figure 1:**
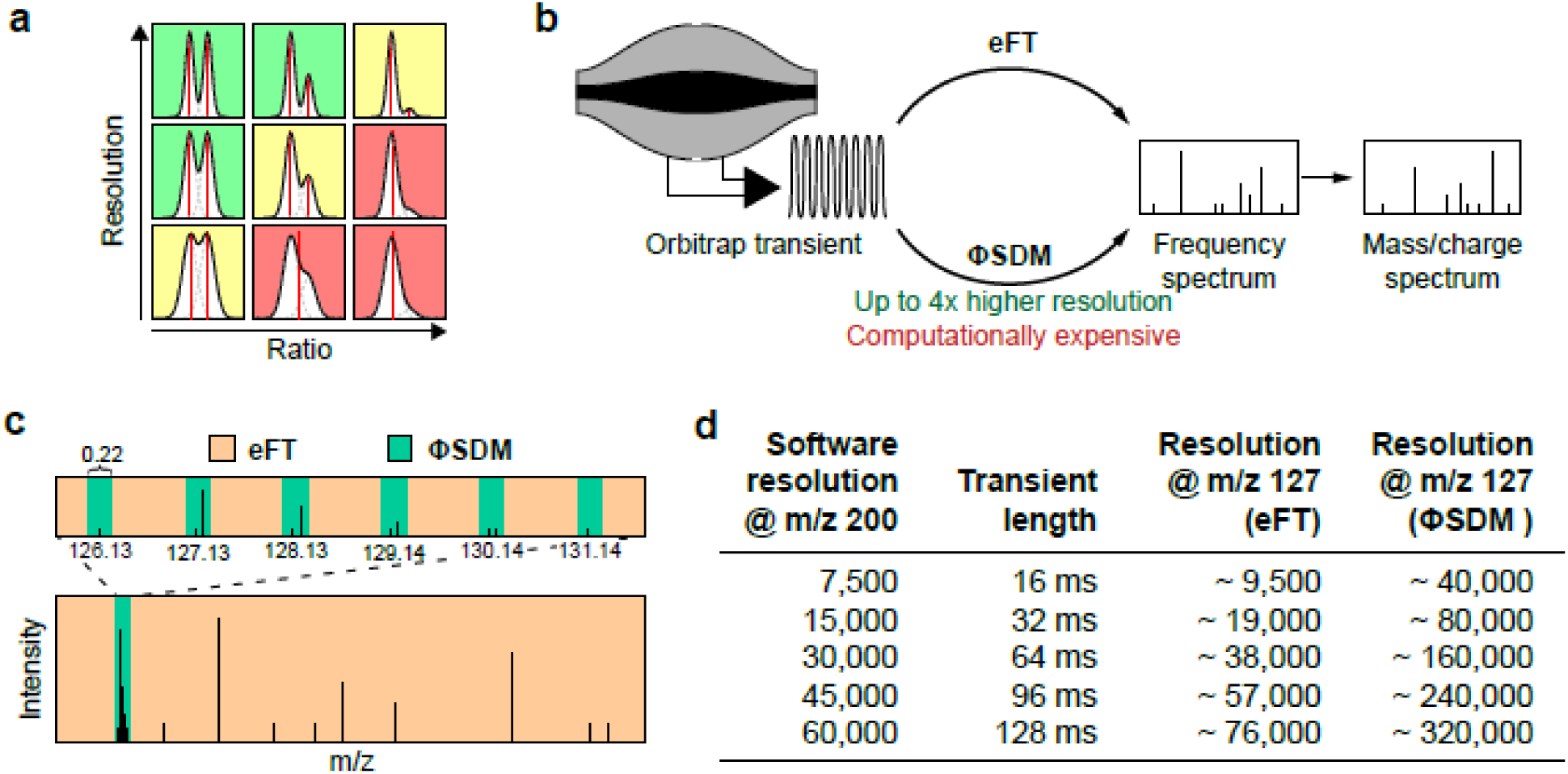
Resolving closely spaced peaks. a) Simulation show two closely spaced Gaussian peaks with different volumes and widths representing different resolutions and intensities. Centroids after basic peak detection is shown in red and the ability of calling peaks resolved is subjectively evaluated with a change in background color where red indicate unlikely to be resolved, yellow is uncertain and green is likely resolved. b) Schematic depiction of the signal processing in Orbitrap mass spectrometry starting from a transient, calculation of the frequency spectrum, followed by conversion to a mass spectrum. c) The employed strategy for limiting ΦSDM to only work on a very small part of the mass spectrum. d) The relationship between resolution settings in the software, the corresponding length of the recorded transient for the ultra-high-field Orbitrap and the resolution in the reporter ion region with either eFT or ΦSDM.

Using the ΦSDM as a super FT resolving approach has recently been shown to increase the mass resolving power of the Orbitrap mass analyzer relative to the eFT method currently implemented on the commercial instrument (Figure 1b).^13^ The ΦSDM algorithm hence supplements the eFT in the data processing of the transient signal to the mass spectrum. Implementation of the ΦSDM algorithm has here been optimized to run on existing instrument hardware with no downsides in acquisition speed (Supplementary Figure S-1a). This was accomplished by limiting the ΦSDM calculations to 6 small m/z ranges, in which the TMT10-plex reporter ions are present (Figure 1c). For the rest of the spectrum, the eFT algorithm was used to enable real-time processing with the current instrument hardware. In greater detail, the promised resolution improvements in this narrow mass range can be seen as substantial (Figure 1d).

To design an empirical test to define the lowest resolution needed to resolve TMT reporter ions with the same integer masses, peptides from a HeLa digest were labeled with the eight central channels from a TMT10-plex kit (Figure 2a). These contain four pairs of closely spaced reporter ions which were mixed in different known ratios. In total, five sample mixtures with known TMT ratios were designed (sample I-V), where four (sample I-IV) contained the four closely spaced peak pairs in ratios from 8:1 to 1:8 (Figure 2a). Sample V contained the more extreme ratios 1:16 and 1:64. All five samples were subsequently measured using data dependent acquisition on short 15 minute gradients with seven different main MS methods. The MS methods used variable resolution settings with either standard eFT or ΦSDM processing for the TMT reporter ion region (Figure 2a).

**Figure 2:**
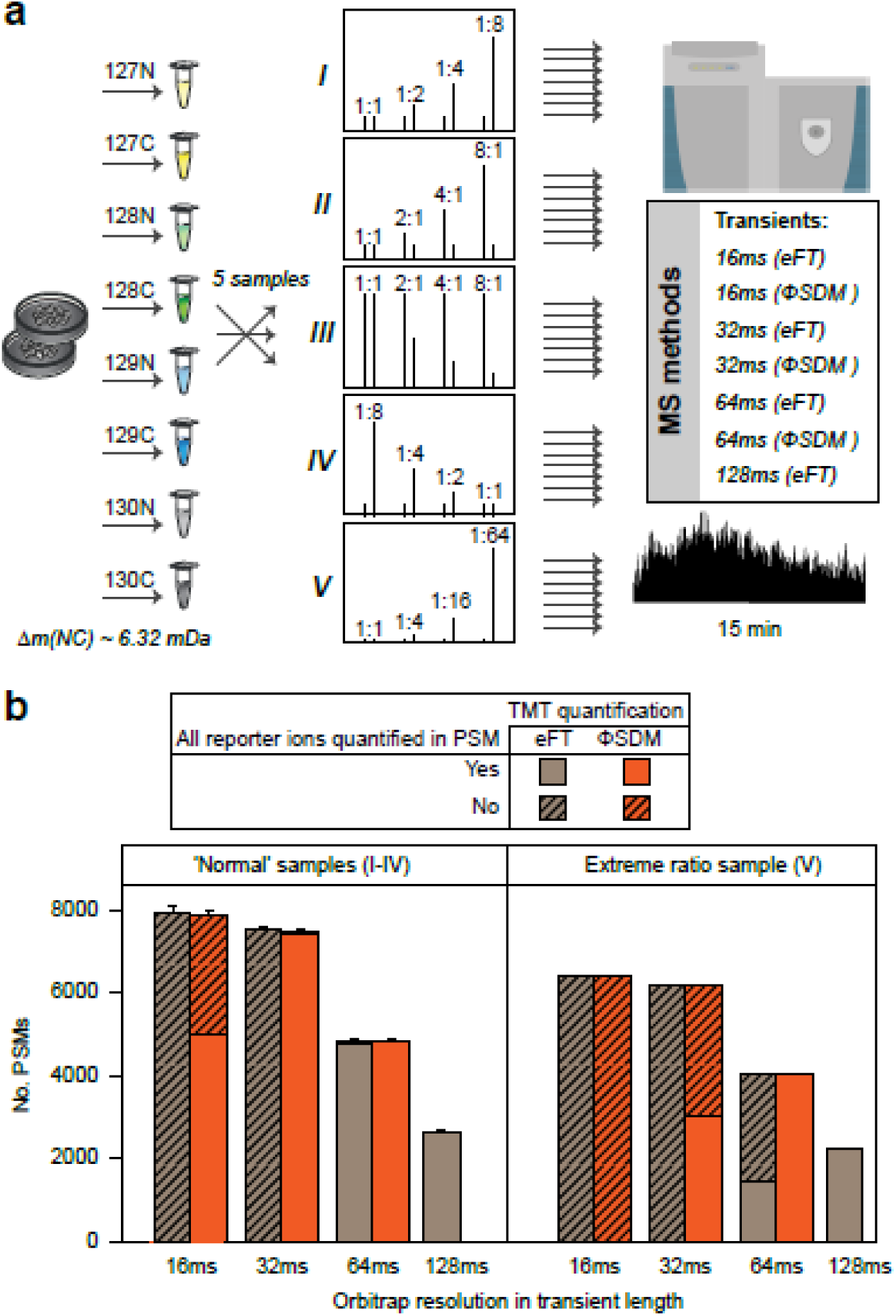
Experimental design and overview results. a) HeLa samples are labeled with the eight central TMT reporter ions, representing four integer mass pairs. These are mixed in five different set of ratios forming five different samples that are measured with seven different MS methods. c) Barchart showing number of identified and quantified spectra for the measured samples. Error bars indicate the standard deviation across different samples.

Samples I to IV were found to behave very similarly with regard to the overall number of identifications and quantifications and they were therefore grouped for this analysis (Figure 2b). The number of peptide spectrum matches (PSMs) increased with short transients; this is a known potential benefit from faster Orbitrap transients when the ion flux is high enough.^19^ A comparison of eFT to ΦSDM showed no significant differences in total number of identifications which confirmed that the computational costs of ΦSDM are kept within the limitations of the current MS hardware.

The challenge with fast acquisition methods is readily apparent when quantified PSMs are investigated (Figure 2b). For eFT-based quantification, the 16 ms and 32 ms transients did not produce any PSMs with all ratios present (fully quantified) indicating problems in resolving the closely spaced peaks. In contrast, the 64 ms eFT method showed great performance for samples I-IV, with >99% of ∼4800 PSMs being fully quantified. The 128ms eFT method, however, leads to 45% less PSMs (∼2600) with similar quantification (>99%) of all PSMs. For sample V, the challenge of quantifying high ratios kicked in and only 35% of PSMs could be fully quantified using the 64 ms eFT method while >99% quantified PSMs was maintained for the 128 ms eFT method that also gave the total highest number of fully quantified PSMs. We found a small software issue that labeled reporter ions as exception peaks in the raw data for the 8:1 ratio with the 128ms eFT transient. This should be fixed in a software update in the future, and we therefore decided to skip this data-point from this part of the analysis.

Comparing eFT to ΦSDM showed no significant differences in the total numbers of identifications, which confirms that the relative computational costs of ΦSDM are practically non-existent as it operates within the limitations of the current MS hardware. Overall, quantitative performance was found improved. The 16 ms ΦSDM methods could fully quantify ∼5000 PSMs from sample I-IV with the 32 ms increasing this to ∼7460 (Figure 2b). With eFT, no reporter ion sets were fully quantified with these very short transient lengths. However, the faster 32 ms ΦSDM method reported here quantified 55% more than the 64 ms method with eFT. Improvements for ΦSDM were also seen for mass accuracies as expected from the introduction (Supplemental Figure S-1b). For the extreme ratio sample V, ΦSDM was able to show an improvement as the 64ms ΦSDM method now kept >99% PSMs fully quantified but with 80% higher total PSMs than the classical 128ms eFT method.

In the analysis so far we considered all quantification ratios as equal, but the relative number of quantified ratios was verified to depend on the ratio (Figure 3a). It is evident that eFT of 16ms transients is not enough to resolve any reporter ion pairs, whereas the 32ms transient has an approximately 70% chance of producing a ratio readout for ratios with changes up to two-fold. The longer 64 ms and 128ms eFT methods generally performed comparably in terms of quantification accuracy and precision with the only difference that the 64ms transient displaying difficulties with the most extreme 1:64 ratio, where only 50% of PSMs were quantified. With ΦSDM replacing eFT for the TMT mass region, the fastest 16ms transient was found to separate ratios up to 4 fairly consistently while higher ratios have >5% chance of producing missing values. Increasing the transient to 32ms was only found to cause problems for the most extreme 1:64 ratio.

**Figure 3:**
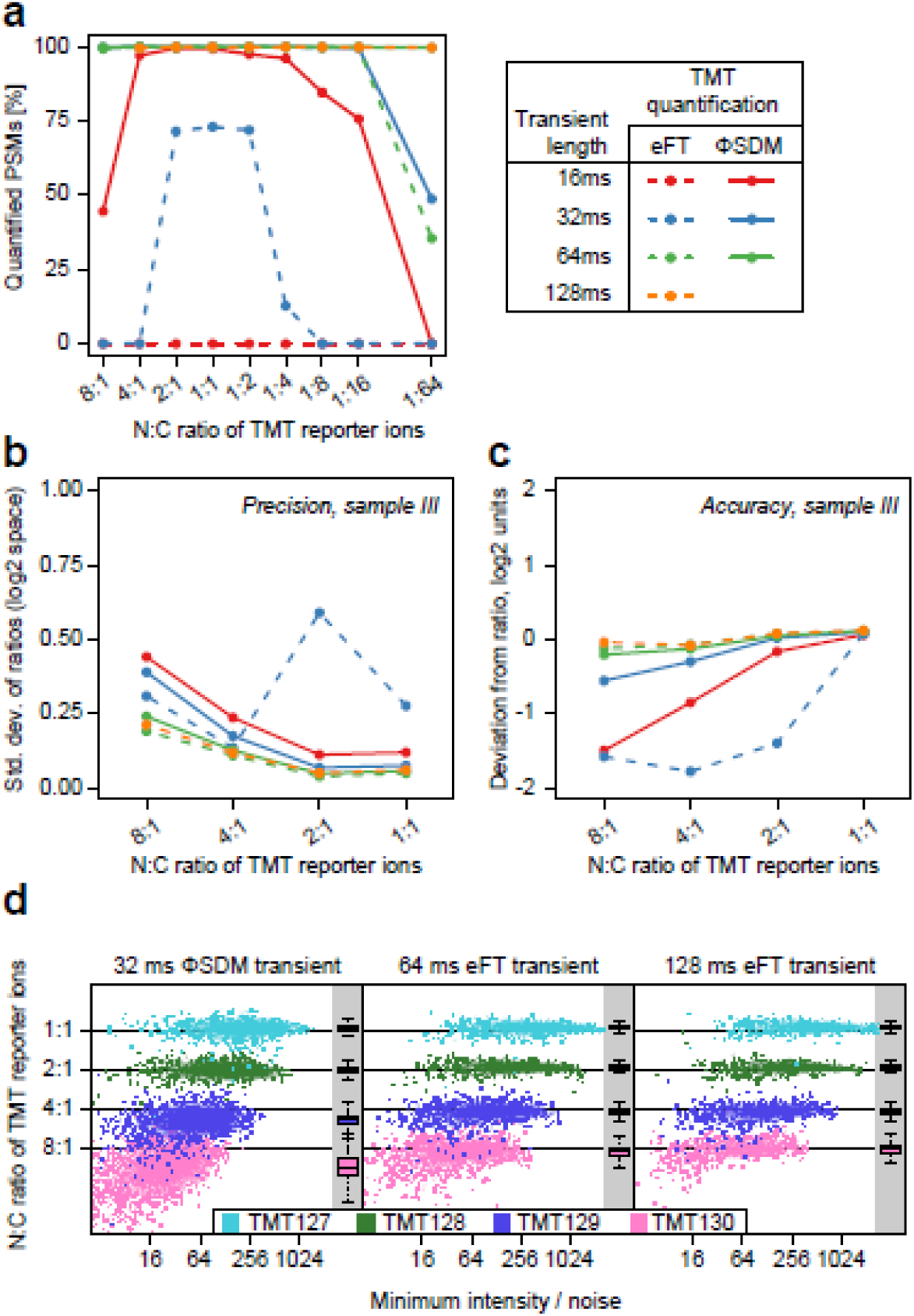
More detailed results. a) Line chart summary of the relative number of quantified PSMs versus the total number of identifications. b) Line chart, using same legend as a, where precision is shown for sample III. c) Line chart, using same legend as a, where accuracy is shown for sample III. d) The measured ratio for four integer masses is shown versus the minimum intensity for sample III. The different plots show the two best performing methods, 32ms + ΦSDM, 64ms + eFT, together with the safe choice, the 128ms + eFT methods with inlaid grey areas showing box plot distributions for the measured ratios. Colored horizontal lines indicate expected ratios.

In more detail, precision in the form of standard deviations of the log transformed ratios was evaluated (example for sample III in Figure 3b with all results in Supplementary Figure S-2a). The 32ms eFT transient, found to give poor qualitative results, gave very variable results in this analysis and can therefore be said to be insufficient for analyzing TMT samples. A general increase in variation for higher absolute ratios was found, highlighting the finding that the precision is best for small absolute ratio differences. The shortest 16ms ΦSDM transient was found to also have the worst precision. ΦSDM and eFT has similar precision at absolute ratios up to 2 but at higher ratios ΦSDM worsened faster than eFT. Accuracy, measured as the deviation from the expected ratio in log2 space, was also investigated (example sample III in Figure 3c with all results in Supplementary Figure S-2a). Results were similar to precision with the two short transients, 32 ms eFT and 16ms ΦSDM, were found to have the worst accuracy. The 32ms ΦSDM showed worse accuracy for absolute ratios above 2 providing more extreme ratios than ground truth. The longer transients perform the best with minimal difference between 64 ms and 128 ms eFT.

The effect of longer transients performing better caused an investigation into signal-to-noise or in this case, intensity-to-noise for the arguably most interesting MS methods (Figure 3d). This showed that intensity-to-noise values were unable to explain the majority of the observed difference between the methods although it can be observed that precision and accuracy worsens for larger absolute ratios at very low intensity-to-noise. As additional controls, the samples were measured with MS method controls which validated that the fill times setting did not affect the outcome of the above mentioned analyses (Supplementary Figure S-2b).

## CONCLUSION

This systematic comparative study shows that two methods in particular seems relevant for future TMT10-plex or even higher multiplexing experiments. The 32ms transient with ΦSDM provided the overall best performance with the highest number of quantified PSMs and lowest costs in precision and accuracy for absolute ratio changes up to 16-fold. The slower, 64ms transient with eFT had slightly better precision and accuracy for larger ratios with the same absolute ratio limitation of 16. This limitation is practically small as absolute ratios above 16 between reporter ions in the same integer mass channel are rarely reported in the literature. However, if larger ratios are expected, longer transients can be used. These findings should be helpful for making decisions on what transient to use for proteomics investigations relying on reporter ion quantification with high resolution requirements. We expect that the 32ms transient with ΦSDM will become the method of choice in the future as ΦSDM accelerates experiments utilizing isobaric reporter ions of TMT.

## ASSOCIATED CONTENT

### Supporting Information

The following files are available free of charge at ACS website http://pubs.acs.org:

Figure S-1: Characteristics of transient lengths and FT processing method.

Figure S-2: Additional data on precision and accuracy.

## Notes

The authors declare the following competing financial interest(s): K.A., A.K., D.G., O.L, D.M., and A.M. are employees of Thermo Fisher Scientific, the manufacturer of the Q Exactive HF-X instrument used in this research.

## ACKNOWLEDGMENT

Work at The Novo Nordisk Foundation Center for Protein Research (CPR) is funded in part by a generous donation from the Novo Nordisk Foundation (grant no. NNF14CC0001). Part of this work has been funded as part of the MSmed project that has received funding from the European Union’s Horizon 2020 Research and innovation program under grant agreement no. 686547. We would like to thank the PRO-MS Danish National Mass Spectrometry Platform for Functional Proteomics and the CPR Mass Spectrometry Platform for instrument support and assistance. J.V.O. was supported by the Danish Cancer Society (R90-A5844 KBVU project grant).

## ABBREVIATIONS

ACN, acetonitrile; CID, collisionally-induced dissociation; eFT, enhanced Fourier transform; FA, formic acid; FT, Fourier transform; Gnd-HCl, guanidine hydrochloride; LC, liquid chromatography; LFQ, label free quantification; m/z, mass-to-charge; MS, mass spectrometry; MS/MS, tandem mass spectrometry; PSMs, peptide spectrum matches; TFA, trifluoroacetic acid; TMT, tandem mass tag; ΦSDM, phase-constrained spectrum deconvolution method

